# Greater gray matter volume in somatosensory and parietal regions in elite skiers compared with other athletes

**DOI:** 10.64898/2026.05.10.724084

**Authors:** Kento Nakagawa, Kazuyuki Kanosue

## Abstract

Elite athletes exhibit sport-specific neural adaptations, yet it remains unclear whether such changes reflect general effects of training or the unique demands of individual sports. Skiing requires postural control and whole-body coordination under dynamically unstable environments, placing high demands on somatosensory processing and sensorimotor integration. The present study aimed to identify structural brain characteristics specific to elite skiers by comparing them with athletes from other sports disciplines and non-athletes. T1-weighted MRI data were analyzed using voxel-based morphometry in 13 skiers, 23 non-ski control athletes and 25 non-athletes. Whole-brain analysis comparing skiers with non-ski athletes revealed a significant cluster showing greater gray matter volume in skiers compared with non-ski athletes in the left postcentral gyrus, extending into the superior parietal lobule. The identified cluster primarily encompassed cytoarchitectonic Areas 2 and 5L. These regions are involved in higher-order somatosensory processing and multisensory integration. Importantly, region-of-interest analysis demonstrated that gray matter volume within this cluster was greater in skiers compared with non-ski athletes and non-athletes, with no difference between non-ski athletes and non-athletes. These findings highlight the relative prominence of structural adaptations within somatosensory–parietal networks, reflecting the unique integration of proprioceptive and other sensory information required for elite skiing. Overall, these findings provide evidence for sport-specific structural brain differences in elite athletes and highlight the importance of somatosensory and parietal regions in sensorimotor integration relevant to skiing. These findings may have implications for understanding neural markers of expertise and may inform future approaches to training and performance evaluation in skiing.

## Introduction

Elite skiers demonstrate exceptional performance under highly unstable environmental conditions, requiring precise control of posture and movement on snow. While previous studies on skiing performance have primarily focused on physiological and biomechanical factors such as muscle volume and strength (1), the neural basis underlying these superior abilities remains poorly understood.

A defining characteristic of skiing, compared to various other sports, is the continuous requirement for postural control under unpredictable and dynamically changing surface conditions. This places strong demands on balance ability and its underlying sensory systems, particularly proprioceptive and vestibular functions. Indeed, previous studies have reported superior balance ability in skiers (1), enhanced postural responses to perturbations (2), and improvements in balance following short-term ski training (3-5). Furthermore, proprioceptive function has been shown to be enhanced in elite skiers compared with controls (6) and to improve across the competitive season (7), suggesting that somatosensory processing plays a critical role in skiing performance.

Neural systems exhibit use-dependent plasticity, and the specific nature of training and environmental demands is known to shape distinct neural adaptations. Previous neuroimaging studies in athletes have demonstrated sport-specific structural and functional changes, particularly in motor-related regions such as the supplementary motor cortex (8, 9) and primary motor cortex (10-12), as well as in the cerebellum (13, 14). In addition, sports requiring advanced spatial processing, such as gymnastics and golf, have been associated with structural differences in the parietal lobule (15, 16). However, most of these studies compared athletes with non-athletes, making it difficult to determine whether observed differences reflect sport-specific adaptations or more general effects of physical training.

To address this limitation, the present study compared elite skiers with athletes from other sports disciplines in addition to non-athletes. This design allows for the identification of neural characteristics that are more specifically associated with the unique demands of skiing rather than general athletic training. Given the critical role of somatosensory processing and multisensory integration in maintaining balance under unstable conditions, we hypothesized that skiers would exhibit plastic structural changes in the primary somatosensory cortex and parietal regions involved in sensorimotor integration and the transformation of sensory information into body-centered representations (17), which are thought to contribute to postural control (18, 19). To test this hypothesis, we employed voxel-based morphometry (VBM) to examine group differences in gray matter volume across the whole brain.

## Materials & Methods

### Participants

Thirteen skiers (ski group), twenty-three athletes from other sports (control athlete group: ConA) and twenty-five non-athletes (NonA) participated in this study. All participants in the ski and ConA groups were elite-level athletes who had competed at the national level or higher. The ski group included athletes from multiple skiing disciplines (e.g., alpine, cross-country, and ski jumping), reflecting the diverse demands of competitive skiing. The ConA group consisted of athletes engaged in sports other than skiing (e.g., baseball, soccer, and track and field). The NonA group did not engage in competitive sports. There were no significant differences between the three groups in age (one-way ANOVA: *F*(2, 58) = 0.865, *p* = 0.427), gender proportion (Pearson’s χ^2^ test: *χ*^*2*^(2) = 0.14, *p* = 0.931). Years of athletic experience also did not significantly differ between the ski and ConA groups (unpaired t-test, *p* = 0.075) (see **Table 1**). In addition, absolute volume of gray matter (*F*(2, 58)) = 0.861, *p* = 0.428) and white matter (*F*(2, 58) = 0.865, *p* = 0.714) did not significantly differ between the three groups. All participants were right-handed, except for one left-handed participant in each group. Written informed consent was obtained from all participants prior to participation. The study was approved by the Human Research Ethics Committee of Waseda University (Ethical Approval Number: 2021–092) and conducted in accordance with the Declaration of Helsinki.

**Table 1.**
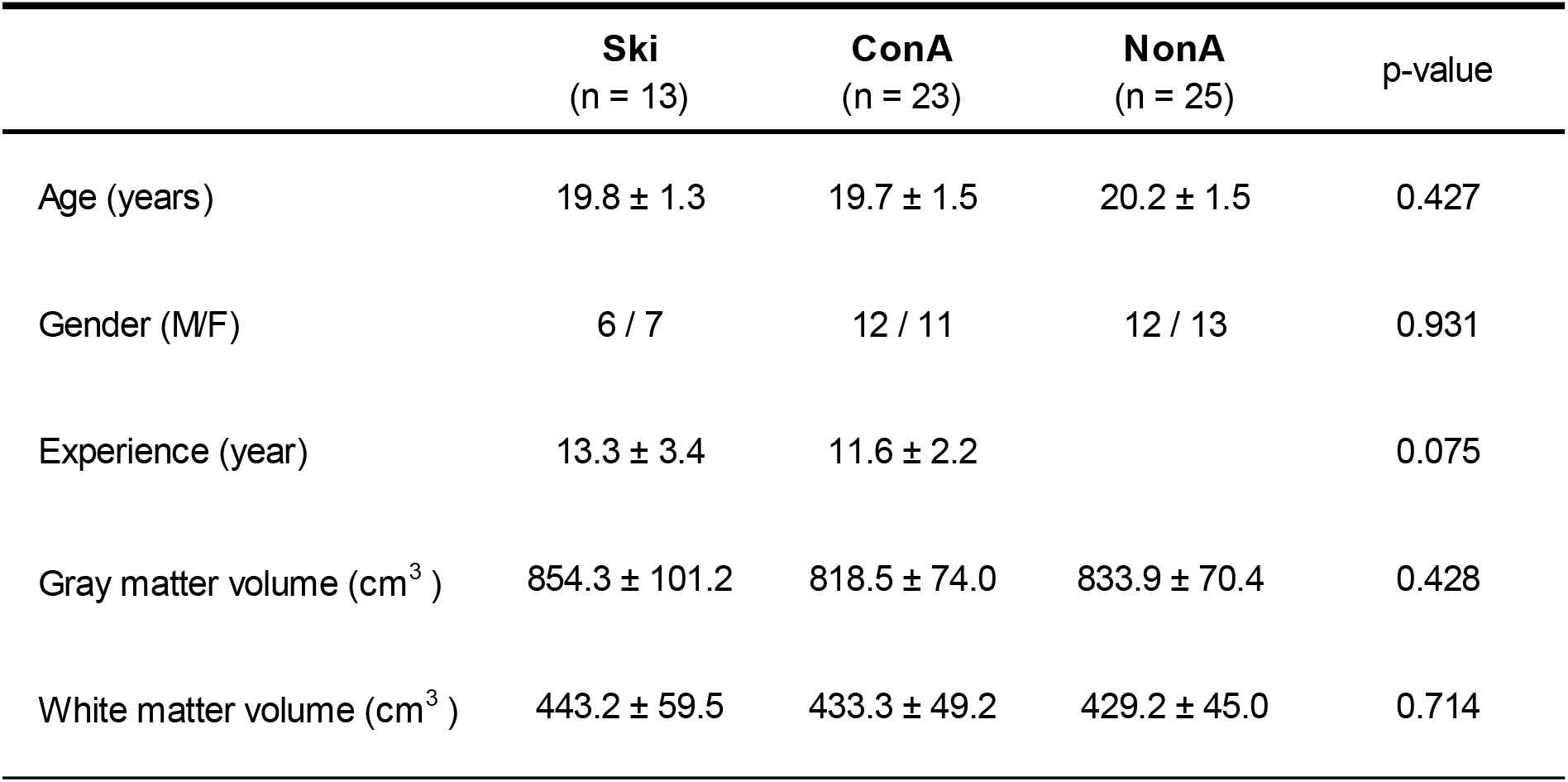
Participant characteristics.

### MRI Acquisition

Structural images were acquired using a 3T MRI scanner (GE SIGNA Premier) with a 48-channel head coil. High-resolution T1-weighted images were obtained using an axial MPRAGE sequence (TR = 2206 ms, TE = 3.4 ms, TI = 900 ms, flip angle = 8°, field of view = 224 × 224 mm^2^, matrix size = 280 × 280, 248 slices, voxel size = 0.8 × 0.8 × 1.6 mm^3^, slice thickness = 1.6 mm).

### Voxel-Based Morphometry (VBM) Analysis

VBM analysis was performed using SPM12 (Wellcome Centre for Human Neuroimaging, London, UK). T1-weighted images were segmented into gray matter, white matter, and cerebrospinal fluid using the standard SPM segmentation procedure based on the ICBM template. Subsequently, gray matter images were spatially normalized to the Montreal Neurological Institute (MNI) space using high-dimensional DARTEL normalization. The normalized gray matter images were modulated to preserve volume information and smoothed with an 8-mm full-width at half-maximum (FWHM) Gaussian kernel.

### Statistical Analysis

Whole-brain VBM analysis was performed to compare gray matter volume between the ski and ConA groups using a voxel-wise two-sample t-test. Age, gender, and total brain volume (calculated as the sum of gray and white matter volumes) were included as covariates to control individual differences in brain size. The statistical threshold was set at a voxel-level threshold of p < 0.001 (uncorrected), with cluster-level correction for multiple comparisons using false discovery rate (FDR) at p < 0.05. Given the exploratory nature of whole-brain VBM analysis, cluster-level FDR correction was applied.

To further examine the pattern of group differences, mean gray matter volume values were extracted from the significant cluster identified in the whole-brain analysis. These values were compared across the three groups (ski, ConA, and NonA) using one-way ANOVA, followed by post hoc pairwise comparisons with Bonferroni correction. These region-of-interest (ROI) analyses were conducted to characterize group differences and should be interpreted with caution due to the non-independence of ROI selection.

## Results

In the whole-brain VBM analysis, a significant cluster showing greater gray matter volume in the ski group compared with the ConA group was identified in the left postcentral gyrus (cluster-level FDR-corrected *p* = 0.032; voxel-level *p* < 0.001, uncorrected), with the cluster extending into the superior parietal lobule based on its spatial extent and cytoarchitectonic labeling (**Table 2** and **Figure 1A**). These areas were predominantly assigned to cytoarchitectonic Areas 2 and 5L based on probabilistic mapping. The peak voxel of this cluster was located at MNI coordinates (−34, −39, 60) with a T value of 5.20. No brain regions showed significantly greater gray matter volume in the ConA group compared with the ski group. In the ROI analysis, based on the cluster identified in the whole-brain comparison, a one-way ANOVA revealed a significant main effect of the group (*F*(2, 58)) = 12.610, *p* < 0.001). Post-hoc analysis revealed that gray matter volume extracted from the significant cluster was greater in the ski group than in the ConA and NonA groups (both *p* < 0.001), with no significant difference between the ConA and NonA groups (*p* = 0.281) (**Figure 1B)**.

**Table 2.** Significant cluster showing greater gray matter volume in skiers than in control athletes (ConA)

| Cluster |  | Peak |  | MNI coordinates<br>(x, y, z) | Macroanatomy | Cytoarchitecture<br>(JuBrain) |
| --- | --- | --- | --- | --- | --- | --- |
| p-value<br>(FDR-corrected) | Cluster size | p-value<br>(uncorrected) | T value |  |  |  |
| 0.032 | 728 | < 0.001 | 5.20 | -34, -39, 60 | Postcentral Gyrus (33.0%)<br>Superior parietal lobule (35.0%) | Area 2 (40.2%<br>Area 5L (17.8%) |

**Figure 1.**
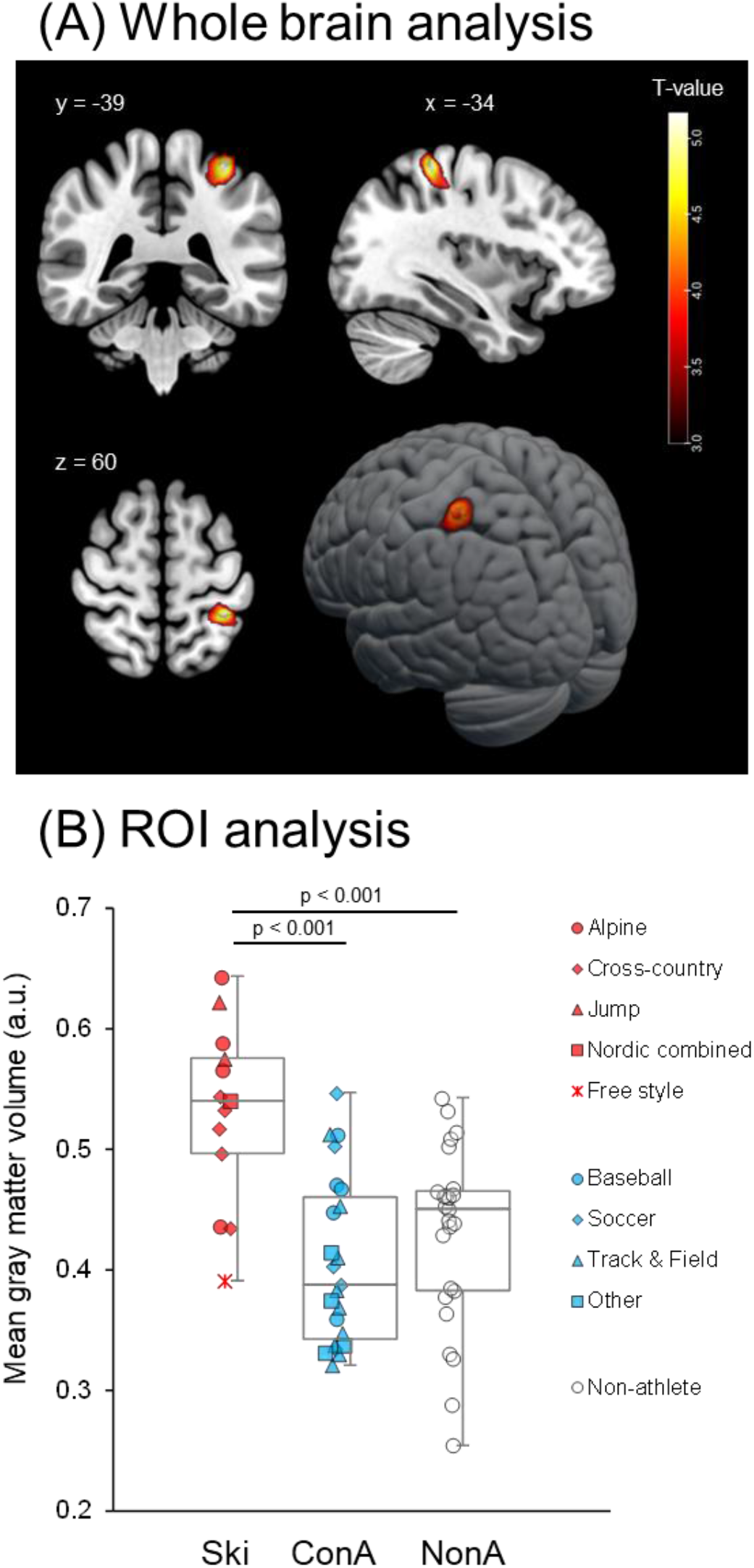
Whole-brain voxel-based morphometry (VBM) results and region-of-interest (ROI) analysis. (A) Brain regions showing significantly greater gray matter volume in the ski group compared with the control athletes (ConA) group (voxel level p < 0.001, uncorrected, cluster-level FDR-corrected p < 0.05). The color bar represents T-values. (B) Individual mean gray matter volume values extracted from the significant cluster (left postcentral gyrus extending into the superior parietal lobule). The ski group showed significantly greater values than both the ConA and non-athlete (NonA) groups (both p < 0.001), with no significant difference between the ConA and NonA groups. Different symbols indicate individual sports disciplines.

## Discussion

### Structural adaptations in somatosensory and parietal regions in skiers and functional significance in skiing performance

The present study investigated sport-specific structural brain differences by comparing elite skiers with athletes from other sports disciplines and non-athletes. This design allowed us to isolate neural characteristics associated with the unique demands of skiing, beyond general effects of athletic training. Using whole-brain VBM analysis, we found that skiers exhibited greater gray matter volume in the left postcentral gyrus extending into the superior parietal lobule. These findings suggest that structural differences in somatosensory and parietal regions may reflect adaptations to the unique sensorimotor demands of skiing, particularly those related to postural control and whole-body coordination under unstable environmental conditions.

A key finding of this study is that the observed structural difference was localized not in motor-related regions but in somatosensory and parietal areas. Previous neuroimaging studies in athletes have frequently reported structural and functional changes in motor-related regions (10, 11, 13, 14). In contrast, the present results highlight a different pattern of adaptation, emphasizing regions involved in sensory processing and integration rather than motor execution per se. This discrepancy may be explained by the use of other elite athletes as the control group. Because both groups had extensive training and motor expertise, structural characteristics of motor-related regions may have been comparable, whereas differences emerged in somatosensory–parietal regions that are more specifically tuned to the demands of skiing, which require continuous adjustment of posture and movement under unstable environmental conditions. Importantly, the inclusion of a non-athlete group in the ROI analysis provides additional support for this interpretation. No significant difference was observed between non-ski athletes and non-athletes, suggesting that the observed structural differences in somatosensory–parietal regions are unlikely to reflect general effects of athletic training. Instead, these findings point toward sport-specific adaptations associated with the unique sensorimotor demands of skiing.

The postcentral gyrus, particularly somatosensory cortex (S1) Area 2, plays a critical role in higher-order somatosensory processing by integrating proprioceptive and tactile inputs to construct representations of body position and movement. This hierarchical organization, in which Area 2 integrates information across multiple body segments, has been well described in primates (20, 21). Consistent with this view, recent electrophysiological studies have demonstrated that neural activity in Area 2 reflects the kinematic state of the whole arm during movement (22). The superior parietal lobule (including Area 5L) is known to function as a key hub for integrating multisensory information from visual and somatosensory systems with motor-related signals, supporting sensorimotor transformations required for coordinated actions and postural control (17). The greater gray matter volume observed in these regions in skiers may therefore reflect enhanced processing and integration of somatosensory information required for maintaining balance and controlling whole-body movements on unstable surfaces. Notably, the peak coordinate of the observed cluster was located between previously reported representations of the hand and lower limb within the primary somatosensory cortex (23, 24). Although VBM does not provide direct evidence of somatotopic organization, this intermediate location likely corresponds to representations of proximal body regions such as the shoulder, upper arm, or trunk. This interpretation is further supported by Mani, Izumi (2), who demonstrated that superior postural control in skiers involves specific coordination of trunk muscles. Such functional demands associated with whole-body stabilization likely drive the structural adaptations observed in the intermediate regions of the somatosensory cortex in the present study. However, this interpretation should be considered with caution due to the indirect nature of structural imaging.

### Methodological considerations and limitations

Several limitations should be acknowledged. First, the sample size was relatively small and unbalanced between groups, which may limit statistical power and generalizability. This limitation partly reflects the difficulty of recruiting highly trained elite athletes with comparable competitive backgrounds. In addition, the extracted gray matter values did not reveal clear clustering according to specific sports disciplines within each group; however, the present sample size was not sufficient to formally evaluate sport-specific sub-group differences. Future studies with larger and more homogeneous samples will be necessary to determine whether distinct sports are associated with more fine-grained structural adaptations. Second, the individual plots (Fig. 1B) revealed considerable variability in GMV even among skiers within the same discipline (e.g., alpine). Whether this inter-individual variance explains specific skiing skills, such as dynamic balance or adaptability to changing snow conditions, remains to be elucidated. Future studies should incorporate quantitative assessments of ski-performance to investigate the direct correlation between brain structure and technical expertise. Third, although cluster-level FDR correction was applied, more conservative correction methods such as family-wise error correction did not yield significant results, and thus the findings should be interpreted with caution. Fourht, the cross-sectional design does not allow causal inferences regarding whether the observed structural differences are the result of long-term training or pre-existing characteristics. Longitudinal studies are needed to clarify the direction of these effects. Finally, MRI data for the skiers were acquired during the off-season, which may have led to an underestimation of the immediate, training-related neural adaptations, as seasonal fluctuations in brain structure and metabolism have been reported in athletes (25), and proprioceptive function has been shown to vary within a season in elite skiers (7). However, the fact that significant structural differences were still observed during this period suggests that these changes do not merely reflect transient, activity-dependent fluctuations. Instead, our findings may reflect stable and consolidated neural adaptations resulting from long-term intensive training. Such persistent structural characteristics in the somatosensory–parietal regions may constitute a foundational neural scaffold for elite skiing performance that remains robust even during the off-season, when the continuous and dynamic sensorimotor challenges inherent to movement on snow are absent.

From a practical perspective, monitoring the structural characteristics of the somatosensory–parietal regions could potentially serve as a supplementary metric for talent identification or evaluating long-term training adaptations. Furthermore, our findings provide a spatial target for neuromodulation strategies. Recent evidence suggests that non-invasive brain stimulation can enhance athletic performance by modulating cortical excitability (26, 27). While the application of brain stimulation in elite sports is still in its infancy, it is a rapidly evolving field. Therefore, identifying the optimal cortical targets is a critical first step. In this context, the identified regions in the present study offer a promising anatomical basis for future neuromodulation-based training aimed at optimizing sensorimotor integration in elite skiers.

In summary, this study demonstrates that elite skiers exhibit greater gray matter volume in somatosensory and parietal regions compared with other athletes. These findings suggest that sport-specific neural adaptations in skiing are characterized by enhanced somatosensory processing and multisensory integration rather than changes limited to motor execution systems.

## Acknowledgements

This study was supported by JSPS KAKENHI (Grant Number 22H03498, 20K19584 and 18H04087).

